# Evaluation of peptidomimetic inhibitors

**DOI:** 10.64898/2025.12.12.693909

**Authors:** Christopher Lenz, Krishna Saxena, Stefan Knapp

## Abstract

Scientific research in drug discovery relies on robust, versatile, and well-established methodologies. One effective strategy for targeting disease-relevant proteins, such as E3 ligases, involves rational drug design using degron peptides as starting points for compound development. Here, we present a comprehensive set of complementary assays for the evaluation of degron-based binders, enabling their integration into an efficient drug discovery pipeline. We introduce screening strategies such as Differential Scanning Fluorimetry (DSF) and Fluorescence Polarization (FP), alongside both solution-based and immobilization-based biophysical techniques, including Isothermal Titration Calorimetry (ITC) and Surface Plasmon Resonance (SPR) for reliable affinity determination. To assess intracellular interactions with full-length target proteins, we also employ Nano Bioluminescence Resonance Energy Transfer (NanoBRET). Together, these methods establish a robust framework for the discovery and characterization of degron-based peptidomimetic compounds.

## 1 Introduction

In the field of drug discovery, the exploration of endogenous mechanisms plays a crucial role in understanding disease-relevant processes. Recently, a novel approach called targeted protein degradation (TPD) has emerged as a promising strategy, aiming to degrade pathogenic proteins rather than merely inhibiting their biological activity. **[1]** PROteolysis TArgeting Chimeras (PROTACs) have emerged as one of the most important pharmacological modalities of TPD strategies. PROTACs have greatly expanded targeting capabilities, extending small molecule-based drug development via the ability to engage disease-causing proteins previously considered “undruggable”, triggering their degradation by the ubiquitin-proteasome system (UPS). **[2]** PROTACs are bifunctional molecules, consisting of an E3 ligase ligand, a linker-moiety and a warhead that binds to the target, which is then recognised as a neo-substrate by the UPS, resulting in selective target polyubiquitination and subsequent degradation. **[3]** An important step in the development of innovative PROTACs is the discovery of small-molecules tightly binding to E3 ligases. **[4]**

Degron sequences act similarly to PROTACs, as they regulate the degradation of certain proteins by targeting them to the UPS. These short motifs however, are either an inherent feature of protein sequences or they can be acquired by post-translational modifications. **[5]** Degrons typically interact directly with E3 ligases, making them ideal candidates for ligase-recruiting elements in PROTAC design. By mimicking the sequence of a degron and synthesizing peptidomimetics based on this motif, attachment points for E3 ligases can be developed. **[6–8]**

For example, the E3 ligase Tripartite Motif Containing 7 (TRIM7), recognizes proteins with a C-terminal glutamine preceded by a hydrophobic residue via its C-terminal PRYSPRY domain, subsequently targeting them for proteolytic degradation (Figure 1). In this context, the following protocols describe five distinct methods and evaluation strategies for the assessment and development of peptidomimetic ligands, featuring exemplary data derived of strategic approaches to target TRIM7. **[6]**

**Figure 1.**
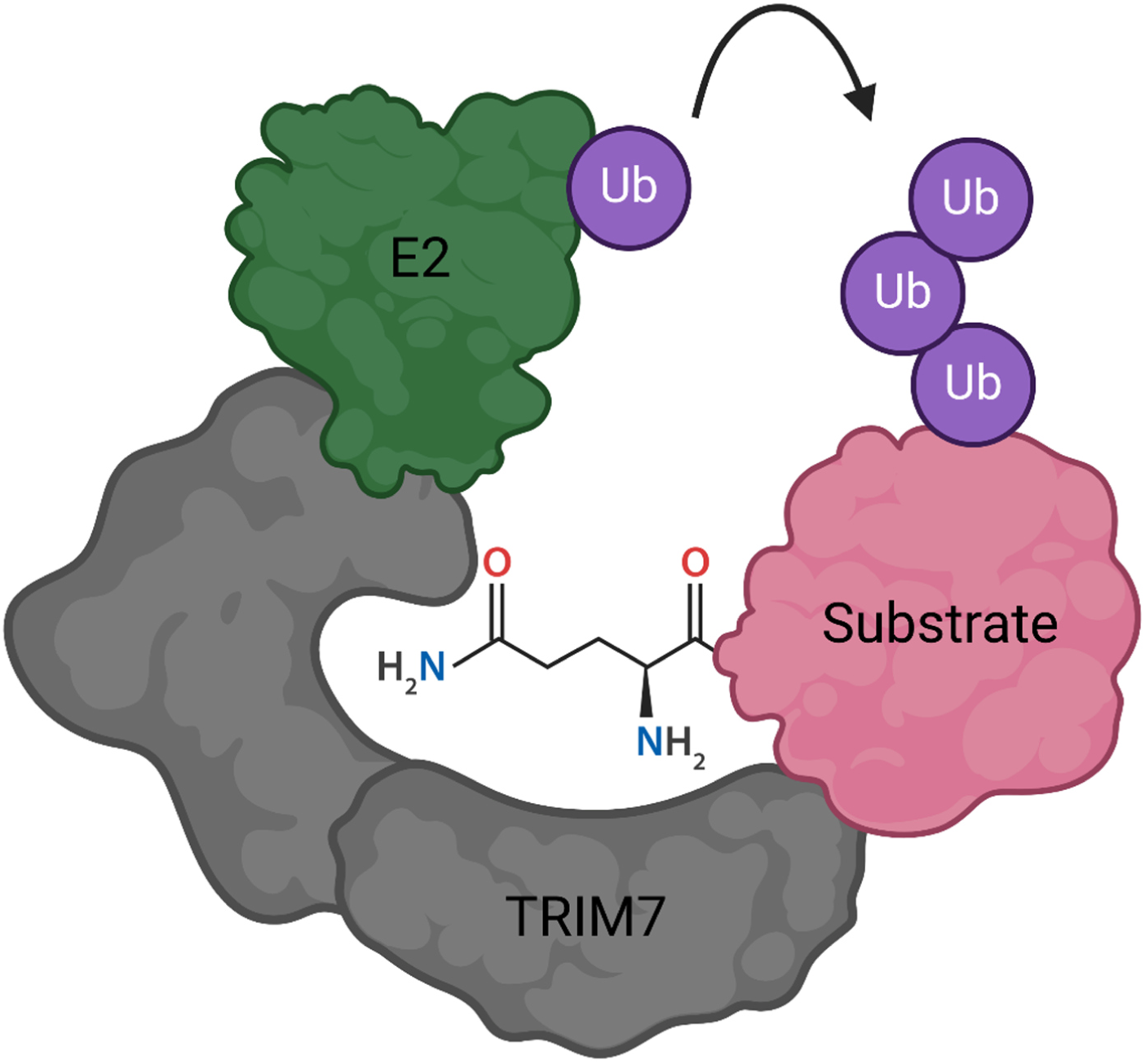
Schematic illustration of TRIM7- and E2-mediated polyubiquitination of a substrate bearing a recognized glutamine residue.

## 2 Materials

In the following sections, a purified E3 ligase protein is required for all *in vitro* protocols. Therefore, it is recommended to either purchase the protein commercially or express and purify it yourself. As a reference, Muñoz Sosa and Lenz et al. **[6]** provided an exemplary description of E3 ligase domain expression and purification.

### 2.1 Differential Scanning Fluorimetry (DSF)

1. DSF buffer: 20 mM HEPES, pH 7.5, 150 mM NaCl, 1 mM TCEP, 5x SYPRO Orange
2. Dimethyl sulfoxide (DMSO)
3. Recombinant E3 ligase protein
4. Peptidomimetic compounds/peptides
5. 384-Well PCR Plate
6. Serological pipettes
7. Pipette controller
8. MCE membrane filter, 0.45 µm pore size
9. Polyolefin sealing film
10. Real-Time PCR instrument
11. Automated liquid dispenser
12. 1.5 mL and 2 mL tubes

### 2.2 Fluorescence Polarization (FP)

1. FP buffer: 50 mM HEPES, pH 7.4, 150 mM NaCl, 1 mM TCEP, 5% Glycerol, 0.05% Tween20
2. Dimethyl sulfoxide (DMSO)
3. Cyanine5 (Cy5)-labelled peptide
4. Recombinant E3 ligase protein
5. Peptidomimetic compounds/peptides
6. Black 384-well flat-bottom plate
7. Serological pipettes
8. Pipette controller
9. MCE membrane filter, 0.45 µm pore size
10. Microplate reader
11. Transcreener-specific FP module (FP 590 675 675)
12. Automated liquid dispenser
13. 1.5 mL and 2 mL tubes

### 2.3 Surface Plasmon Resonance (SPR)

1. Running buffer: 10 mM HEPES, pH7.5, 150 mM NaCl, 1 mM TCEP, 0.05% Tween20
2. Protein dilution buffer: Acetate pH 4.5-5.5; BisTris pH 6.0-6.5
3. Recombinant E3 ligase protein
4. Peptidomimetic compounds/peptides
5. Dimethyl sulfoxide (DMSO)
6. 483 mM N-ethyl-N′-(3-(dimethylamino)propyl)carbodiimide (EDC)
7. 10 mM N-hydroxysuccinimide (NHS)
8. 1 M ethanolamine
9. 1 M NaCl in 50 mM NaOH
10. PP microplate, 96-well, U-shape
11. 25 mL polystyrene reservoirs
12. Serological pipettes
13. Pipette controller
14. MCE membrane filter, 0.2 µm pore size
15. SPR instrument
16. Sensor chip CM5
17. Sensor chip NA
18. Sensor chip SA
19. 1.5 mL and 2 mL tubes
20. 15 mL tubes
21. 7 mm polypropylene plastic vials
22. Rubber caps, type 3
23. 15 mm polypropylene plastic vials
24. Rubber caps, type 5

### 2.4 Isothermal Titration Calorimetry (ITC)

1. ITC Buffer: 20 mM HEPES, pH 7.5, 150 mM NaCl, 1 mM TCEP
2. Dimethyl sulfoxide (DMSO)
3. Contrad 70
4. Recombinant E3 ligase protein
5. Peptidomimetic compounds/peptides
6. Serological pipettes
7. Pipette controller
8. MCE membrane filter, 0.45 µm pore size
9. 15 mL ultra-centrifugal filter, 10 kDa cut-off
10. ITC instrument
11. 1.5 mL and 2 mL tubes
12. 15 mL tubes

### 2.5 Nano Bioluminescence Resonance Energy Transfer (NanoBRET)

1. Dulbecco’s modified eagle’s medium (DMEM), 10% fetal bovine serum (FBS), 1% penicillin-streptomycin (PS)
2. Opti-MEM I reduced serum medium
3. Dulbecco’s phosphate-buffered saline (DPBS)
4. 0.05% Trypsin-EDTA
5. Dimethyl sulfoxide (DMSO)
6. Eukaryotic expression vector with full length protein fused to Nluc (Nluc-E3-ligase DNA) **[9]**
7. Transfection Carrier DNA
8. BODIPY-labelled peptide/peptidomimetic
9. Peptidomimetic compounds/peptides
10. HEK293T cells or different appropriate cell line
11. Nano-Glo Substrate (Promega)
12. FuGENE HD Transfection Reagent (Promega)
13. White 384-well flat-bottom plate
14. Microplate reader with a luminescence filter pair (450 nm BP filter and 610 nm LP filter)
15. Automated liquid dispenser
16. Serological pipettes
17. Pipette controller
18. Cell culture flask T75
19. 1.5 mL and 2 mL tubes
20. 15 mL tubes
21. Automated cell counter
22. CO_2_ incubator
23. Laminar flow hood

## 3 Methods

In general, the use of an automated liquid dispenser, such as the Echo 550 acoustic dispenser (Labcyte) is highly recommended to allow a fast throughput of measurements in a significantly reduced assay volume, resulting in much lower cost per data point.

### 3.1 Differential Scanning Fluorimetry

Differential Scanning Fluorimetry (DSF) is an easy to use, but broadly applicable high-throughput *in vitro* screening method to determine and analyse the apparent melting temperature (Tm) of a purified protein in bound and unbound state. **[10]** As the temperature increases, the protein unfolds, exposing its hydrophobic regions, which can be bound by a dye such as SYPRO Orange. This interaction causes a substantial increase in fluorescence, serving as an indicator of the protein unfolding transition (*see* **Note 1**). **[11]**

Samples used in DSF are prepared as triplicates to determine the standard deviation for each compound analysed. A known binder should be added as a positive control (e.g. peptide degron sequence) as well as a negative/no compound control with only the protein and solvent present to reference data and determine the relative ΔTm. Moreover, an additional protein-free control for auto-fluorescent compounds or buffer ingredients are helpful to prevent the misinterpretation of artefactual, protein-independent fluorescence. For the DSF measurements conducted in this work, we used a QuantStudio 5 Real-Time PCR machine, the Applied Biosystems QuantStudio Design & Analysis Software and Applied Biosystems Protein Thermal Shift Software.

#### 3.1.1 DSF sample preparation and data generation

1. Dilute the E3 ligase protein to 10 µM in 0.45 µm filtered DSF Buffer (*see* **Note 2**)
2. Transfer 10 µL of diluted protein to each well of a 384-well PCR Plate
3. Add peptides/peptidomimetic compounds to reach 100 µM (10x protein concentration) (*see* **Note 3**) with 1-5% DMSO using an automated liquid dispenser to three separate wells for triplicates, additionally leave a triplicate as an “only protein” control (*see* **Note 4**)
4. Seal the plate using a polyolefin sealing film and place it into the Real-Time PCR instrument
5. Setup the experiment method by setting the cover temperature to 105°C, the starting temperature to 25°C, the temperature increase-rate to 0.05°C/s and the final temperature to 85°C
6. Select respective wells with your samples and define ligand (protein) and analyte (peptides/peptidomimetics)
7. Start the run
8. After ∼25 min, evaluate the data using a Protein thermal shift analysis software

#### 3.1.2 DSF data evaluation

For the evaluation of your data, it is important to note that there are two most commonly used approaches.

1. ΔTm-Derivative analysis
2. ΔTm-Boltzmann fit

Both evaluation methods can be used to determine Tm-values, however the strategy used for assessment of the data should stay the same.

For the derivative analysis, the derivative of the fluorescence signal with respect to the temperature (d(Fluorescence)/dT) (Figure 2B) is being determined. Here, the Tm is corresponding to the maximum reached for the derivative.

**Figure 2.**
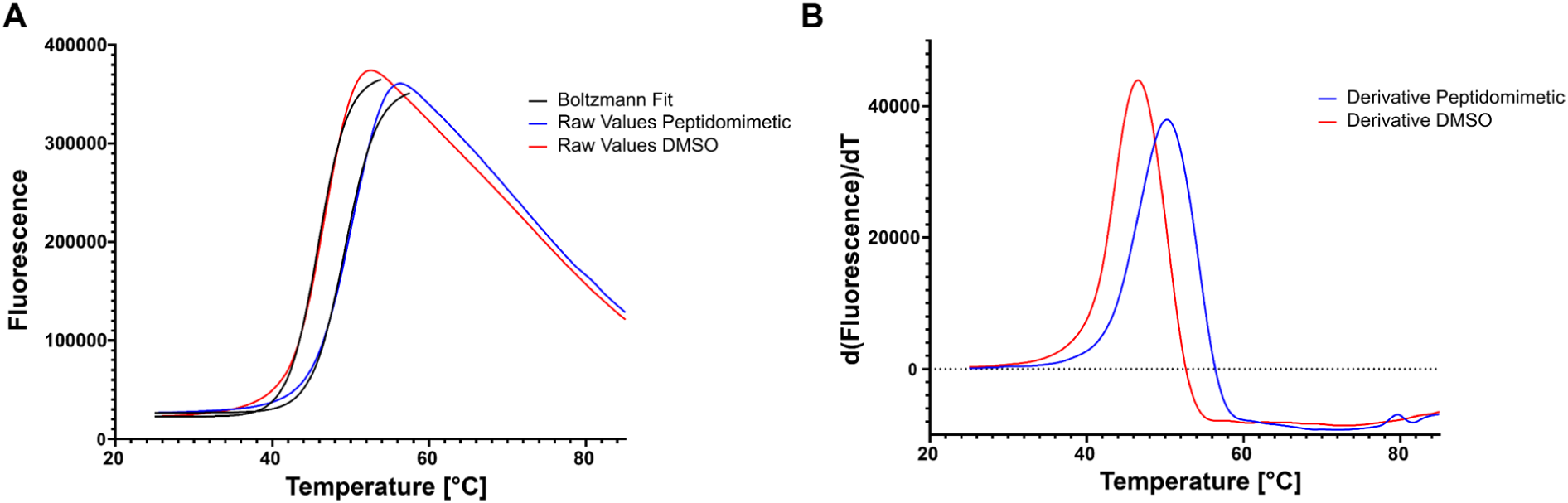
(A, left) Exemplary thermal shift melting curves of TRIM7^PRYSPRY^ with DMSO (blue) and TRIM7^PRYSPRY^ with a peptidomimetic compound (red) as well as the overlaid Boltzmann-fit (black). (B, right) Derivative curves of the same data as in (A).

The Boltzmann fit (Figure 2A) applies a sigmoidal model to fit the melting curve as a function of temperature. Respective Tm values are determined as the temperature at which 50% of the protein is unfolded, corresponding to the inflection point of the curve.

By providing an “only protein” reference, ΔTm-values can be determined via:

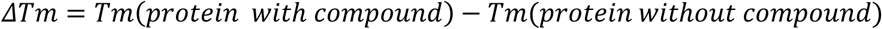

### 3.2 Fluorescence Polarization

The fluorescence polarization (FP) assay is an advanced technique to monitor molecular interactions in solution and is based on the principle that small fluorescently labelled molecules (typically <1.5 kDa) exhibit slower rotational motion when bound to a larger partner (typically >10 kDa). Polarization values, calculated as the ratio of the difference between vertically and horizontally polarized light to their sum, serve as indicators of the ligand state, with low polarization values corresponding to the unbound state and high values indicating the bound state. **[12]**

This method is suitable for screening molecules, such as peptidomimetic compounds against E3 ligase targets, enabling an initial assessment of ligand affinity. The FP assay format also serves to confirm the binding results of investigated compounds determined by DSF. In general, Tm values correlate with binding affinities observed in an FP competition-based approach but the magnitude of ΔTm shifts may depend on the E3 ligase used. By conjugating a fluorescent dye, such as Cy5, to the relevant degron peptide sequence, a tracer molecule can be easily generated (*see* **Note 5**). Prior to its use in a competition assay format, the tracer molecule needs to be analysed via tracer titration with the protein (E3 ligase) to determine its effective affinity.

All samples are prepared as triplicates, additionally if the assay is performed for the first time, different tracer concentrations should be used to determine optimal polarization intensities. For the following FP measurements, we used a Pherastar FSX plate reader.

#### 3.2.1 FP tracer titration assay

1. Dilute the tracer in 0.45 µm filtered FP buffer to reach different tracer concentrations (e.g. 0.1-0.3x expected K_D_ of the tracer) (*see* **Note 6**)
2. Add up to e.g. 100 µM E3 ligase to 100 µL of each tracer suspension
3. Serially dilute the protein-tracer suspension using a 1:1 ratio by following these steps: Sequentially add 50 µL of tracer-only suspension to 50 µL of the preceding E3-ligase-tracer suspension. Repeat this process for a total of 10-15 dilution steps.
4. Transfer 3x 10 µL of each dilution to three separate wells of a black 384-well flat-bottom plate
5. 3x 10 µL of the initial tracer suspension in FP buffer with no protein should also be added to the plate to set the baseline (unbound tracer sample)
6. Incubate the plate at room temperature (RT) in dark conditions for 40 min
7. Read the perpendicular and parallel intensities using a microplate reader
8. Calculate the mean polarization values [mP] for each concentration:

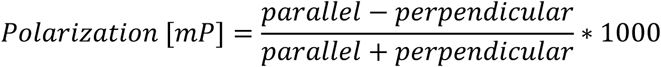

1. 9. Calculate the baseline-corrected polarization (P) by subtracting the polarization value of the unbound tracer sample from all other sample values. Plot the corrected polarization values P against the corresponding tracer concentrations (C) (Figure 3A). Perform a nonlinear regression fit to determine the K_D_ value of the tracer molecule:

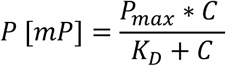

P is the polarization at a given concentration, P_max_ represents the maximum polarization, C is the concentration of the tracer, and K_D_ the dissociation constant of the tracer

**Figure 3.**
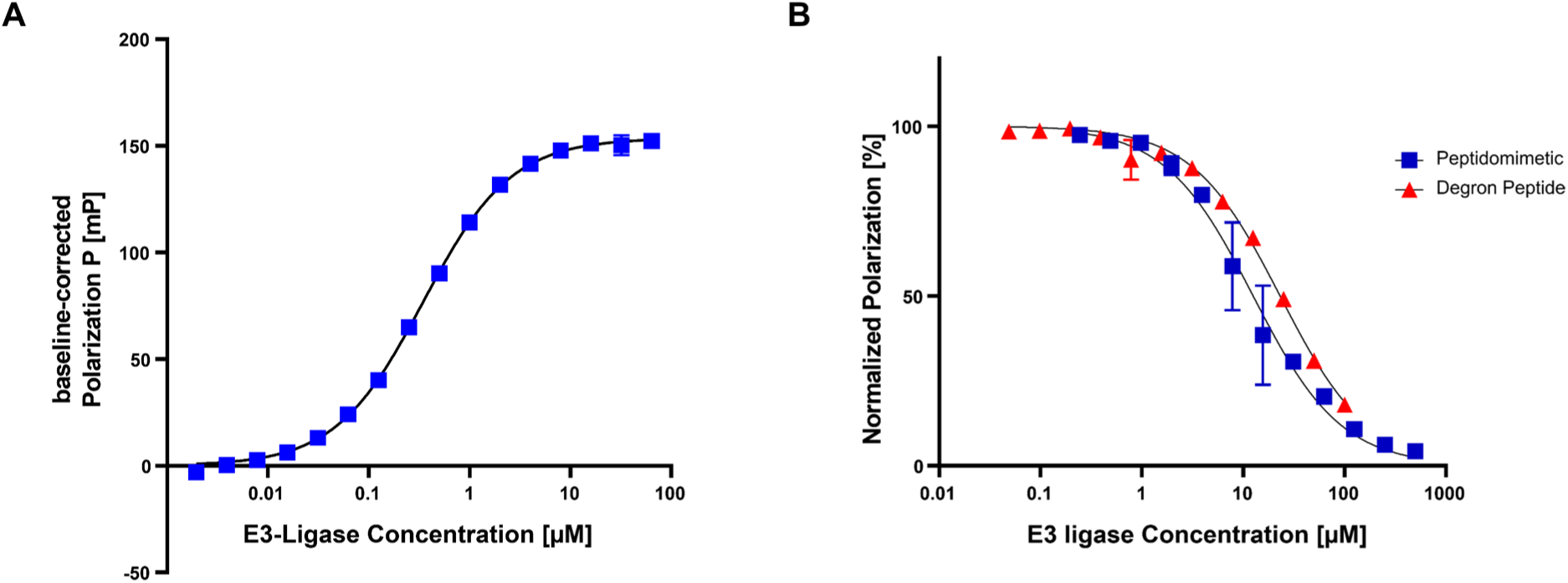
(A, left) Exemplary FP tracer titration curve for the interaction of E3 ligase TRIM7^PRYSPRY^ to a peptidomimetic tracer. (B, right) Dose response competition curves of a peptidomimetic compound (blue) and the degron peptide (red) against TRIM7^PRYSPRY^. Non-linear regression fits are overlaid in black.

#### 3.2.2 FP dose response competition assay

1. Prepare the tracer solution by diluting the tracer in FP buffer to a final concentration of 0.1-0.3x K_D_ of the tracer-protein interaction, as determined previously (*see* **3.2.1**)
2. Before adding the protein to the tracer solution, take out 100 µL to have an unbound tracer sample
3. Add protein at a concentration of 2-3x K_D_ to generate the protein-tracer mixture, then take out 100 µL for the 100% bound tracer sample
4. Transfer 10 µL to each well of a black 384-well flat-bottom plate
5. Use an automated liquid dispenser to titrate different concentrations of peptidomimetic compounds at 1-5% final DMSO concentration to each well of the protein-tracer mixture (*see* **Note 7**)

Alternatively, if you don’t have an automated liquid dispenser:

1. Prepare the tracer solutions diluted in FP buffer as described before (*see* **3.2.2 steps 1-2**)
2. Set aside 100 µL of the protein-tracer mixture for each compound and add the desired final DMSO concentration (1-5%) to the remaining protein-tracer solution
3. Add the highest concentration of compound to the 100 µL protein-tracer mixture set aside, and adjust the final DMSO concentration to the desired level
4. Serially dilute the compound-solution from **step 3** by transferring 50 µL of this mixture to 50 µL of the solution without compound
5. Transfer 10 µL of each concentration into the corresponding wells of a black 384-well flat-bottom plate
6. Incubate the plate for 40 min at RT under dark conditions
7. Measure the perpendicular and parallel intensities using a microplate reader and calculate the mean polarization values (*see* **3.2.1 step 8**)
8. Normalize mean polarization values for each concentration:

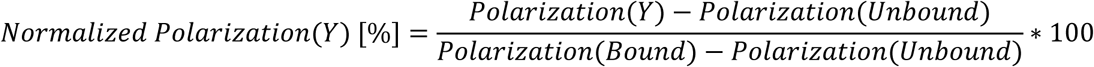

1. Plot the normalized polarization values (Y) vs concentration (C) and perform a nonlinear regression fit to determine the IC_50_ for each peptidomimetic compound (Figure 3B):

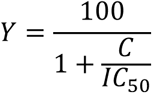

1. To determine K_i_ values for respective compounds using the FP competition assay, the following equation published by Nikolovska-Coleska et al. **[13]** is recommended:

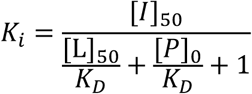

[I]_50_ represents the concentration of free compound at 50% binding, [L]_50_ denotes the free tracer concentration at 50% binding. [P]_0_ corresponds to the free protein concentration at 0% binding, and K_D_ refers to the previously determined dissociation constant of the protein-tracer interaction. For more details, please refer to Nikolovska-Coleska et al. publication.

### 3.3 Surface Plasmon Resonance

Surface Plasmon Resonance (SPR) acts as a label-free, non-fluorescent and versatile analytical technique with high sensitivity, widely used for studying molecular interactions in real-time. The principle of SPR depends on the interaction of polarized light with a thin metal film, typically gold, under conditions of total internal reflection. **[14]** When light hits the metal surface at a specific angle, it excites collective oscillations of conduction electrons, known as surface plasmons. **[15]** These oscillations produce evanescent waves that penetrate a short distance into the adjacent medium. The intensity of these waves is influenced by the refractive index near the chip’s surface. At a specific angle, maximum energy transfer occurs, leading to a noticeable dip in the reflected light intensity. **[14, 16]**

The resonance angle is highly sensitive to changes in the refractive index, caused by interactions on the metal surface. Ligands, such as E3 ligases can be immobilized on the sensor surface, allowing the quantitative detection of binding events with analytes, such as peptidomimetic compounds. The magnitude of the refractive index change directly correlates with the mass and density of the interacting molecules, enabling the determination of kinetic parameters, such as association (k_a_) and dissociation rates (k_d_), as well as the equilibrium binding constant (K_D_). **[14, 16]** To validate potential binders identified in previous screening assays, SPR is an ideal method to accurately determine their respective affinities in a high-throughput manner.

The following protocol is specified for running assays on a Biacore T200 instrument with the Biacore T200 Control/Evaluation Software, using the PRYSPRY domain of the E3 ligase TRIM7, but it is broadly applicable to any other SPR instrument or protein-ligand system. There is a wide variety of immobilization strategies to robustly attach a protein to the surface of a sensor chip, however the two most common approaches, amine coupling on CM5-Chips and biotin-neutravidin (NA)/-streptavidin (SA) coupling will be described.

#### 3.3.1 Preparation steps for immobilization of the protein (*see* Note 8)

1. Prepare 1.5 L running buffer using deionized water and filter with a 0.2 µm pore size membrane filter
2. Take out 500 mL as immobilization buffer and 50 mL running buffer for dilution purposes (*see* **Note 9**)
3. Attach the immobilization buffer to the SPR machine and prime the system
4. Eject the previous sensor chip from the instrument and inject a newly opened sensor chip CM5 or NA-/SA-Chip, then prime again
5. Add 19.4 mL of 100% DMSO to 950 mL running buffer to reach 2% DMSO (for 5% DMSO add 50 mL)

#### 3.3.2 Scouting for an optimal pH level of the protein dilution buffer (for CM5 chips)

1. Start a manual run at 10 µL/min flow-rate on any flow-channel (FC) except for FC1
2. Dilute the protein to 10 µg/mL in Acetate/BisTris buffer pH 4.5-6.5
3. Inject each protein dilution for 30 s and monitor the binding to the chip surface
4. Select the most suitable buffer by choosing the highest pH which is still showing a high response increase for further covalent immobilization procedure (*see* **Note 10**)

#### 3.3.3 Protein immobilization on sensor chip CM5

1. Prepare 100 µL 483 mM N-ethyl-N′-(3-(dimethylamino)propyl)carbodiimide (EDC), 100 µL 10 mM N-hydroxysuccinimide (NHS) and 140 µL 1 M Ethanolamine separately in a 7 mm plastic vial
2. Add up to 700 µL of 10 µg/mL E3 ligase diluted in protein dilution buffer to another 7 mm plastic vial and keep an additional 7 mm plastic vial empty for the mixture of EDC and NHS
3. Select a new immobilization method and choose “Amine coupling” on the 2^nd^ FC with the following settings:
4. Mix and inject 483 mM EDC + 10 mM NHS (1:1) for 420 s at 10 µL/min
5. Inject 10 µg/mL protein diluted in protein dilution buffer at 10 µL/min with a contact time according to the desired immobilization level (*see* **Note 11**)
6. Inject 1 M Ethanolamine for 420 s at 10 µL/min
7. Start the immobilization run and repeat **steps 1-3** for the remaining 3^rd^ and 4^th^ FCs to generate replicate protein surfaces
8. Repeat **steps 1-3** additionally for the reference FC1, but select “Blank immobilization”, leaving out the protein injection

#### 3.3.4 Protein immobilization on a sensor chip NA/SA

1. Start a manual run and select all FCs at 10 µL/min
2. Consecutively inject 1 M NaCl in 50 mM NaOH for 1 min x3 over all four FCs
3. End the run and start a new manual run by selecting FC2 at 10 µL/min
4. Inject 10 µg/mL biotinylated E3 ligase diluted in immobilization buffer until the desired protein immobilization level is reached (*see* **Note 11**)
5. Stop the injection and check if the protein-level stays stable
6. Stop the run and repeat **steps 3-5** for FC3 and FC4 to create replicates

#### 3.3.5 Analysis of the dose-response binding of peptidomimetic compounds

1. Design a new method by setting the detection to all four channels with FC1 being the reference channel and the sample compartment temperature set to 25°C
2. Multi cycle kinetics should be used as the experimental strategy with a flow-rate of 30 µL/min as well as at least 60 s contact time and 150 s dissociation time
3. Add the following assay steps while selecting ten compound concentrations, typically ranging from 0.1x to 10x the expected K_D_ for each analyte and include two blanks (running buffer without compound) per analyte. Ten startup injections should be added to ensure a stable baseline, while control sample injections are used to confirm protein integrity over time, using the peptide as a positive control (Figure 4)
4. For a 2% DMSO solvent correction, prepare 10 mL 1% DMSO running buffer and 10 mL 3% DMSO running buffer, using the running buffer that was set aside in **3.3.1 step 2** (*see* **Note 12**)
5. Mix both solutions using the pipetting scheme in Table 1 and transfer each of the 8 mixtures to 7 mm plastic vials
6. Prepare 300 µL of the highest compound concentrations at 2% DMSO using the running buffer that was set aside in **3.3.1 step 2**
7. Transfer the entire volume to a designated well of a 96-well microplate, along with 150 µL of 2% DMSO running buffer into 11 adjacent wells for each compound.
8. Prepare a 1:1 serial dilution by transferring 150 µL from the highest concentration well to the adjacent well and repeat the process nine times sequentially, leaving two samples as blanks.
9. Prepare the control sample with a peptide concentration equal to its K_D_ and transfer the sample to a 7 mm plastic vial
10. Transfer all necessary solutions as well as the 96-well microplate to the SPR metal-rack and inject the rack to the sample compartment
11. Attach the running buffer with 2% DMSO to the SPR instrument, prime the system and eventually start the run
12. After the run has finished, assess the data and plot resulting Sensorgrams (Figure 5A) using the SPR Evaluation Software
13. If steady state was reached for all concentrations, evaluation can be done using the steady state affinity model (Figure 5B) or manually via GraphPad Prism with the following equation:

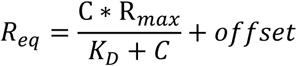

C stands for each concentration of the respective analyte, R_max_ is the maximum response, R_eq_ represents the response at steady state and offset is the response at zero compound concentration

1. 14. For kinetic evaluation of the data (*see* **Note 13**), the 1:1 Langmuir interaction fit can be applied to determine the association rate constant k_a_ and dissociation rate constant k_d_. The affinity can be calculated using the following equation:

**Figure 4.**
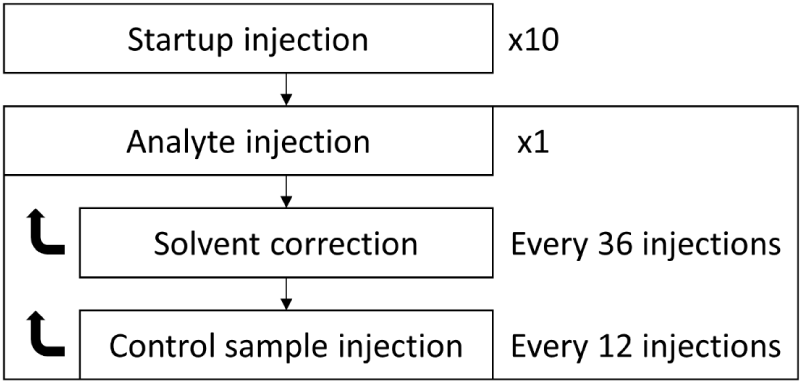
Schematic representation of a general SPR workflow, starting with 10 startup injections, followed by repeated analyte injections, periodically followed by solvent corrections and control sample injections.

**Figure 5.**
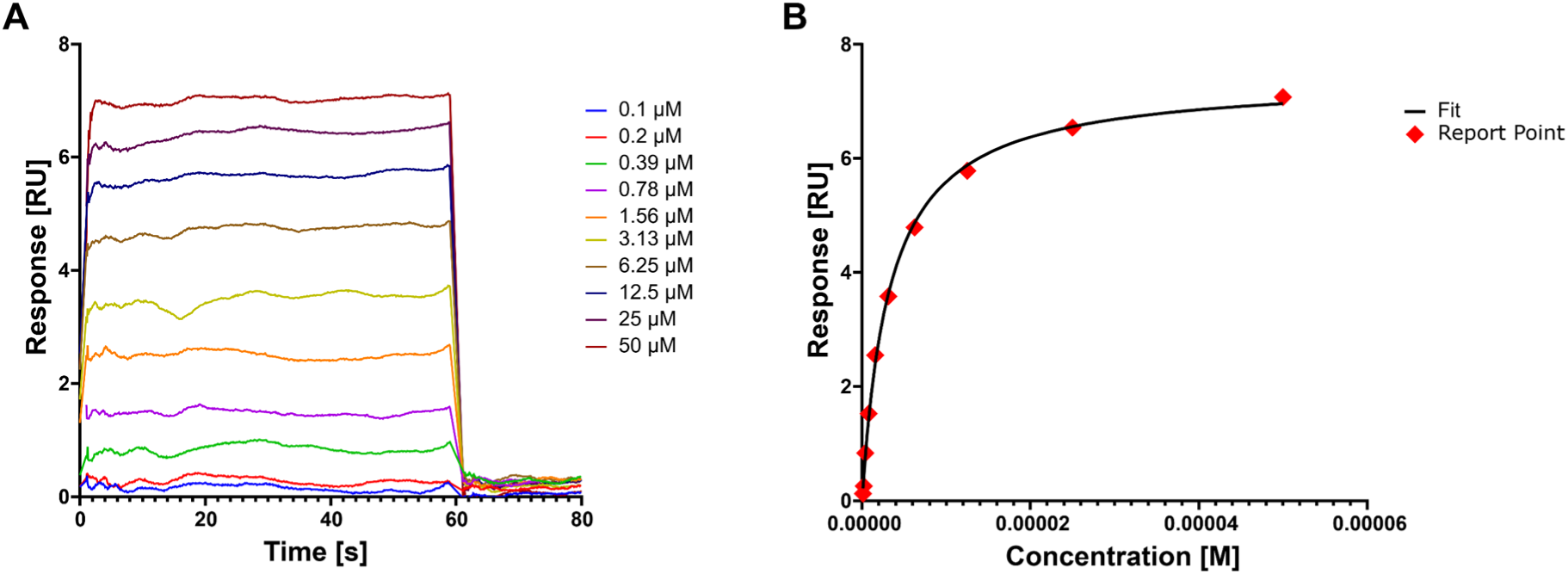
(A, left) Exemplary SPR sensorgram overlay plot of a peptidomimetic compound interacting with TRIM7^PRYSPRY^. (B, right) Steady state affinity analysis of the same data as in (A) with respective report points (red) and overlaid steady state affinity fit (black).

**Table 1.**
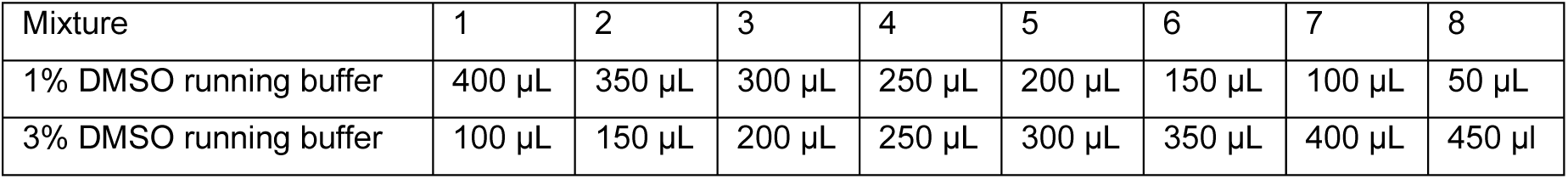
Pipetting scheme for the preparation of the SPR solvent correction.

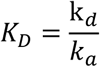

### 3.4 Isothermal Titration Calorimetry

Isothermal Titration Calorimetry (ITC) is another widely accepted and powerful technique in the field of biophysical drug discovery, providing critical insights into molecular interactions. It serves as a complementary method to SPR, as ITC does not rely on binding kinetics but instead determines the thermodynamic profile of interactions. **[17]** While the throughput of this method is quite limited because of its substantial reagent requirements, ITC can be used as a versatile alternative technique to confirm potential binders. By directly measuring the heat released or absorbed during a binding event, ITC enables the determination of key binding parameters, including the binding affinity (K_D_), enthalpy change (ΔH), entropy change (ΔS), and stoichiometry (n) of the interaction. **[18]**

Thermodynamic parameters are calculated using the following equations, while the stoichiometry is derived from the molar ratio of the binding partners in the titration curve:

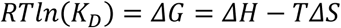

Here, R represents the gas constant, T is the temperature and ΔG is the Gibbs free energy change.

The principle of ITC is based on successive titrations of an analyte (e.g. peptidomimetics) into a sample cell containing its binding partner (e.g. E3 ligase), while the whole process is performed under controlled isothermal and isobaric conditions. **[19]** Each binding event leads to a detectable heat change, which is recorded and analysed to generate a thermodynamic binding profile. **[20]** Unlike most other biophysical techniques, ITC requires no immobilization or modification of interacting molecules, making it a label-free and solution-based method that consistently maintains the native state of both binding-partners. **[21]**

The following protocol is specifically optimized to run ITC for the PRYSPRY domain of E3 ligase TRIM7 on a TA Instruments Nano ITC device. However, it can be applied to any other E3 ligase/protein and ITC instrument by adjusting variable parameters like stirring speed, temperature, buffer conditions, the number of injections, analyte and protein concentrations as well as the DMSO concentration.

#### 3.4.1 Sample preparation

1. Given the high level of sensitivity in ITC, the composition of all components of the ITC buffer need to be identical
2. If necessary, exchange buffer for the protein, using a suitable centrifugal filter with a 10 kDa pore size at 4000 g, adding freshly prepared 0.45 µm filtered ITC buffer at least four times
3. The protein sample (titrand) should have a final concentration of 30 µM in 1.6-1.8 mL ITC buffer with 0.6% DMSO
4. Add 1.8 µL of 50 mM peptidomimetic compound in 100% DMSO to 298.2 µL ITC buffer for 300 µM titrant compound solution

#### 3.4.2 ITC titration experiment

1. Prior to compound titration, all samples need to be degassed to prevent the formation of bubbles which could affect thermal signals
2. Simultaneously, clean the sample cell with detergent (e.g. Contrad 70), followed by rinsing with at least 300 mL deionized water, ensuring the cell is left empty
3. Overfill the sample cell slightly above the desired level with an appropriate amount of titrand, then add 250 µL of the titrant to the syringe, ensuring no bubbles are present.
4. Place the syringe in the sample cell and start stirring at 300 rpm to equilibrate the system until the curve has reached steady state
5. The titration is set to begin with a 4 µL injection of the titrant, succeeded by 27 injections of 8 µL at 25°C

#### 3.4.3 ITC data analysis

1. Analysis is performed by reviewing the thermogram, plotting raw heat rate (µcal/s) vs time (s) and checking for inconsistencies like baseline drifts, noise or injection artefacts (Figure 6 top panel)
2. Integrate the area under each peak to obtain the heat change (ΔQ) for every injection
3. Exclude injection outliers and subtract the baseline ΔQ to remove any noise or systematic deviations
4. Normalize the heat values by dividing the baseline-corrected heat change per injection by the number of moles of injected compound
5. Plot the normalized heat (kcal/mol of injection) against the molar ratio of compound and protein to generate a binding isotherm and confirm that the data follows a sigmoidal curve, indicating saturation of the absolute heat change (Figure 6 lower panel)
6. Select an appropriate fitting model, in this instance a one-site (1:1) binding model is assumed for a single binding site with the following equation:

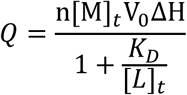

Here, [M]_t_ stands for the total protein concentration, V_0_ represents the total volume of the ITC sample cell and [L]_t_ is the total compound concentration in solution.

**Figure 6.**
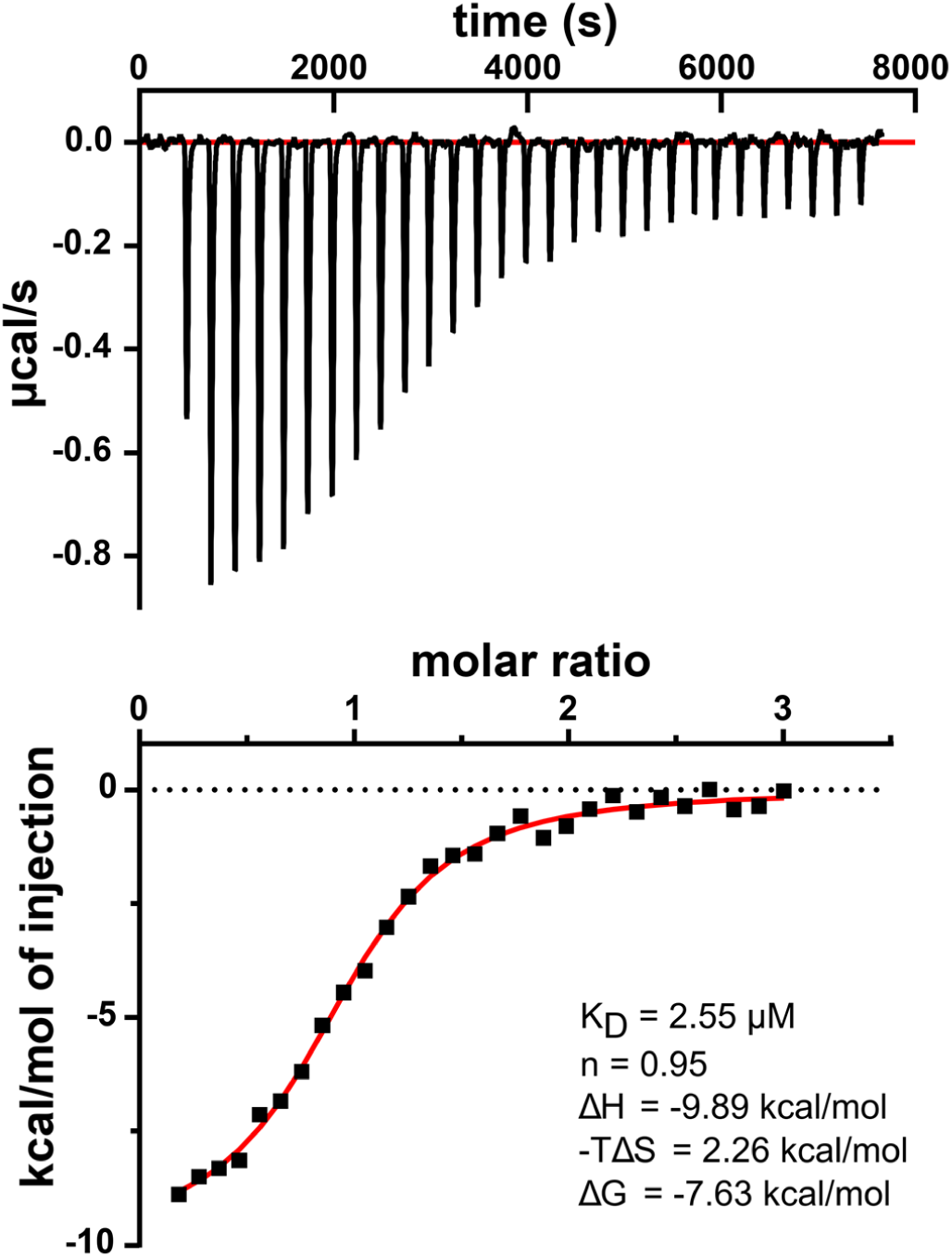
ITC data of a peptidomimetic compound titrated to TRIM7^PRYSPRY^ depicted as a thermogram (top) and integrated heat data (bottom) fitted with a one-site binding model (red). Exemplary results for thermodynamic parameters are shown in the lower panel.

### 3.5 Nano Bioluminescence Resonance Energy Transfer

Cellular assays play a crucial role in confirming potential binders *in vivo*. Here, the focus lies on permeability of the compounds and their engagement with the full-length target in a cellular environment that is difficult to determine with most conventional *in vitro* assays. Nano bioluminescence resonance energy transfer (NanoBRET) is a powerful method which uses proximity induced energy transfer of a bioluminescent enzyme to fluorescent acceptor molecules, to measure this interaction upon substrate consumption. **[22]** In this context, a NanoLuc luciferase (Nluc) acts as the respective enzyme, conjugated N- or C-terminally to an E3 ligase (e.g. TRIM7), exciting a fluorophore (tracer) in vicinity. The tracer molecule needs to be localized in the area of the potential peptidomimetic binding site to facilitate screening based on a competitive manner. **[23]** Transiently transfected human cells like HEK293T are typically being used to study cellular engagement, while the tracer molecule can be based on the peptide degron sequence or a peptidomimetic conjugated to fluorescent dye BODIPY.

The following workflow describes a general NanoBRET assay process used to determine the apparent K_D_ of a tracer molecule to its protein target as well as the IC_50_ of compounds displacing the tracer. Both assay options require transient transfection of the Nluc-tagged E3 ligase before measurement and can be evaluated in intact or permeabilized cellular states. All samples should be prepared as triplicates to ensure reproducibility. Additionally, compounds, tracer, DMSO and digitonin should be dispensed using an automated liquid dispenser to enable rapid and precise distribution. We used a Pherastar FSX plate reader for the following NanoBRET measurements.

#### 3.5.1 Transfection of Nluc-constructs

1. Prepare 500 µL transfection mix, by adding 15 µL FuGENE HD transfection reagent to a mixture of 9 ng/µL Transfection Carrier DNA and 1 ng/µL Nluc-E3-ligase DNA construct in Opti-MEM medium (For the selection and design of functional Nluc-constructs refer to Schwalm et al. **[9]**)
2. Incubate the transfection mix for 15 min at RT
3. Seed HEK293T cells to a final density of 2.0 x 10^5^ cells/mL in DMEM with 10% FBS and 1% PS, then add 500 µL transfection mix to 10 mL cell-solution in a cell culture flask T75
4. Incubate cells over-night at 37°C and 5% CO_2_

#### 3.5.2 NanoBRET tracer titration assay

1. Remove DMEM culture medium and displace with Opti-MEM medium using serological pipettes (*see* **Note 14**)
2. Adjust cell density to 2.0 x 10^5^ cells/mL and transfer 10 µL of transfected cells to each well of a white 384-well flat-bottom plate
3. Titrate different concentrations of the tracer into respective wells with a final DMSO concentration of maximum 1% (*see* **Note 15**)
4. Incubate the plate at 37°C and 5% CO_2_ for 2 h
5. Add 5 µL Nano-Glo substrate diluted 1:200 in Opti-MEM to each well
6. Measure BRET ratio using a microplate reader with a luminescence filter pair [450 nm BP filter and 610 nm LP filter]
7. Lyse cells by adding 25 nL of 20 mg/mL digitonin to each well, then incubate for 5 min
8. Measure BRET ratio again with the same settings as in **step 6** for permeabilized cells

#### 3.5.3 NanoBRET tracer displacement assay

1. Remove DMEM culture medium and displace with Opti-MEM medium using serological pipettes
2. Adjust cell density to 2.0 x 10^5^ cells/mL and transfer 10 µL of transfected cells to each well of a white 384-well flat-bottom plate
3. Dispense tracer at a concentration not exceeding 0.1x K_D_ to each well, as determined in section **3.5.2** (*see* **Note 16**)
4. Add increasing concentrations of peptidomimetic compounds to respective wells at a maximum DMSO concentration of 1% (*see* **Note 17**)
5. Add 5 µL Nano-Glo substrate diluted 1:200 in Opti-MEM to each well
6. Measure BRET ratio using a microplate reader with a luminescence filter pair [450 nm BP filter and 610 nm LP filter]
7. Lyse cells by adding 25 nL of 20 mg/mL digitonin to each well, then incubate for 5 min
8. Measure BRET ratio again with the same settings as in **step 6** for permeabilized cells

#### 3.5.4 NanoBRET Data analysis

The BRET ratio is calculated using the following equation:

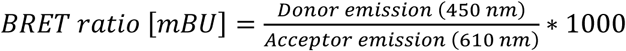

To determine the EC_50_ value of the tracer, the concentration is plotted against the baseline corrected BRET ratio and fitted with a non-linear model (Figure 7A):

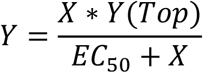

**Figure 7.**
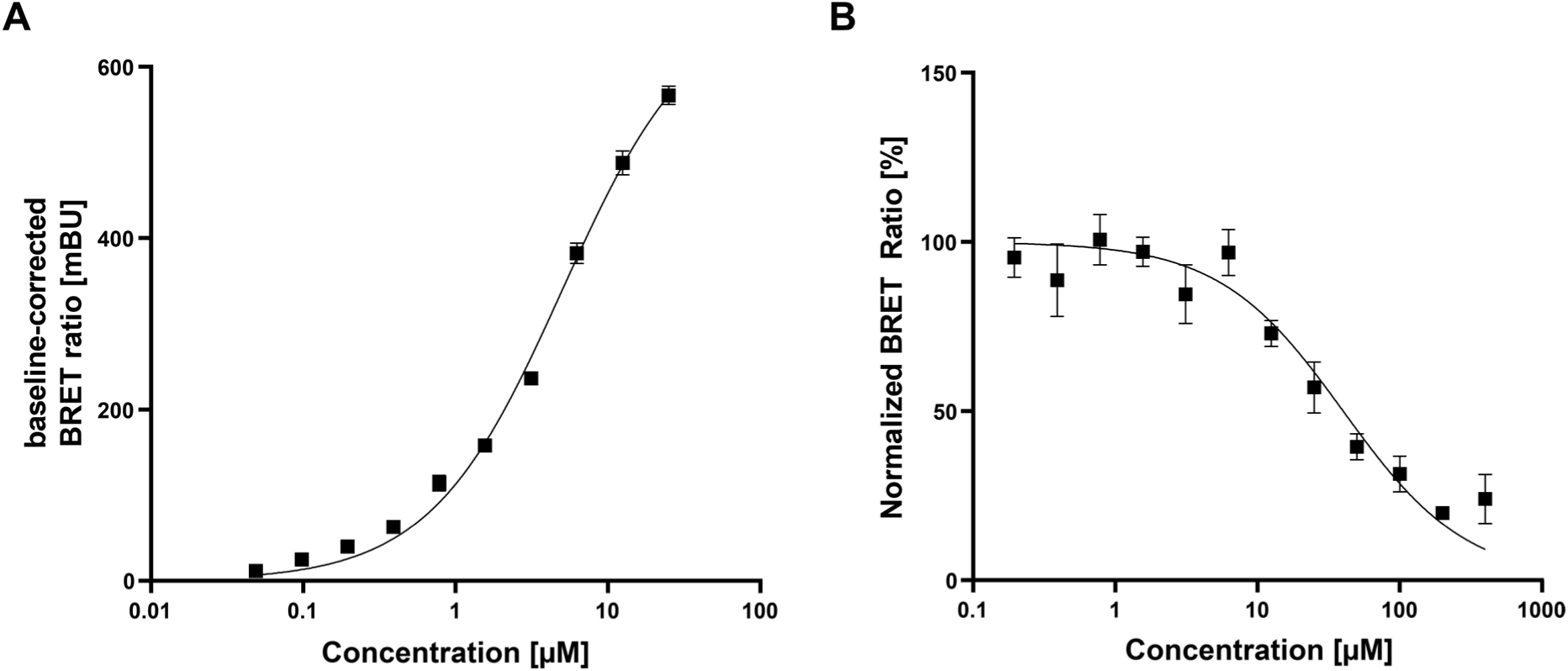
(A) NanoBRET tracer titration of a BODIPY-conjugated peptidomimetic tracer in permeabilized HEK293T cells overexpressing full-length Nluc-TRIM7. (B) Competitive dose-response analysis showing displacement of the tracer molecule by the peptidomimetic compound in permeabilized cells. The respective non-linear regression fit is shown in black as overlaid curves.

Y represents the BRET ratio, X denotes the tracer concentration and Y(Top) corresponds to the BRET ratio at the highest compound concentration.

Displacement data is evaluated using normalized competition data with the following equation to determine IC_50_ values for each compound (Figure 7B):

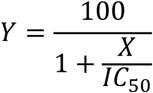

Here, X represents the concentration of the compound and Y is the normalized BRET ratio.

## 4 Notes

1. DSF should not be confused with NanoDSF as they utilise different strategies. NanoDSF monitors unfolding through tryptophan fluorescence, while DSF detects protein unfolding using selective dye fluorescence. **[24]**
2. It is important to get a good fluorescent signal to determine a reliable Tm-value for any E3 ligase. Varying the protein concentration (∼1-20 µM) can help in finding a suitable fluorescence intensity for the screening of compounds with different binding affinities.
3. Try to determine the solubility of any peptidomimetic compound/peptide prior to the measurement to avoid precipitation effects.
4. DMSO concentration should be the same for all compounds and added to the “only protein” control.
5. Even though Fluorescein isothiocyanate (FITC) and dyes with similar spectral properties are among the most widely used fluorophores, red dyes such as Cy5 have gained traction due to the improved performance of modern plate readers. **[25]**
6. If the K_D_ of the peptide is already known, start with a tracer concentration lower than the K_D_.
7. Concentrations of the compounds should cover at least 0.1x to 10x K_D_ of the compound-protein interaction, to ensure enough curvature and the generation of valid plateaus for both maximal and minimal displacement.
8. Every solution and buffer as well as the sensor chip should be warmed up to RT prior to starting the assay to avoid baseline drifts and artefacts.
9. The buffer used to dilute samples needs to be exactly the same as the running buffer to avoid response deviations caused by differences in buffer composition.
10. By using a very low pH, you might risk a loss of protein integrity due to a sharp pH drop.
11. Protein levels should be selected according to reaching an expected maximum response (R_max_) for interacting peptidomimetics of ∼10-20 RU. Calculations can be done via:

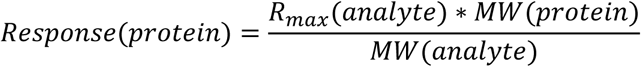

1. For different DMSO concentrations, adjust both solutions to ± 1% relative to the desired final DMSO concentration.
2. For functional kinetic evaluation, the association phase should exhibit visible curvature, while the dissociation phase should show at least 5% dissociation of the compound. **[26]**
3. Opti-MEM medium should be free of PS and FBS to ensure optimal screening conditions, minimizing unspecific interactions as well as high background fluorescence/luminescence signals.
4. Tracer concentrations should be chosen according to the expected K_D_ of its binding motif to cover 0.1x to 10x K_D_. Moreover, a no-tracer control should be added for baseline-correction.
5. For normalization purposes, include a no-tracer control and a sample containing the tracer but without the compound.
6. Select a broad concentration range of compound (0.1x to 10x the expected K_D_) to ensure protein saturation and complete displacement of the tracer.

## Acknowledgements

The authors are grateful for support by the Structural Genomics Consortium (SGC), a registered charity (no. 1097737) that receives funds from Bayer AG, Boehringer Ingelheim, Bristol Myers Squibb, Genentech, Genome Canada through Ontario Genomics Institute [OGI-196], EU/EFPIA/OICR/McGill/KTH/Diamond Innovative Medicines Initiative 2 Joint Undertaking [EUbOPEN grant 875510], Janssen, Merck KGaA (aka EMD in Canada and U.S.), Pfizer, and Takeda. SK would like to acknowledge funding from the German Cancer Consortium (DKTK) at the German Cancer Research Center (DKFZ).

